# It’s raining species: Rainwash eDNA metabarcoding as a minimally invasive method to assess tree canopy invertebrate diversity

**DOI:** 10.1101/2022.03.24.485661

**Authors:** Till-Hendrik Macher, Robin Schütz, Thomas Hörren, Arne J. Beermann, Florian Leese

## Abstract

1. Forest canopies are a highly diverse ecosystem, but despite several decades of intense research, there remain substantial gaps in our knowledge of their biodiversity and ecological interactions. One fundamental challenge in canopy research is the limited accessibility of the ecosystem. Consequently, previous studies have relied on the application of either highly invasive methods such as chemical knockdown, or on time-consuming and expensive setups such as canopy walkways or cranes. Therefore, time- and cost-efficient, ideally minimally invasive yet comprehensive applications are required to help close this knowledge gap. High-throughput metabarcoding of environmental DNA (eDNA) collected from water, soil, or air provides a minimally invasive method for biodiversity assessment, yet its potential for canopy biodiversity monitoring has not been explored.
2. Herein, we conducted metabarcoding of eDNA washed off the canopy via rainwater to explore its monitoring potential. We placed four 1 m^2^ rain samplers beneath the canopies of four different tree taxa prior to a major rain event, filtered eDNA from the collected rainwater, and performed cytochrome c oxidase subunit I (COI) metabarcoding to profile the invertebrate community. Additionally, we collected and identified all specimens present in the rainwater for verification.
3. We detected 50 invertebrate species by eDNA metabarcoding, of which 43 were not physically present in the water sample, thus likely representing true canopy biodiversity signals. Furthermore, we observed distinct species occurrence patterns corresponding to the four tree taxa, suggesting that ecological patterns such as host specificity can be assessed using the method.
4. In conclusion, our study provides a proof of principle that rainwash eDNA metabarcoding offers a minimally invasive and comprehensive method for tree canopy diversity monitoring.

## Introduction

The forest canopy is a particularly species-rich zone. However, despite many decades of intense canopy research, there remain substantial gaps in our knowledge of the biodiversity and ecological interactions of canopy communities (Nakamura et al., 2017). This includes both tropical rainforests with their vast array of undescribed species (Basset et al., 2012), and temperate forests (Sallé et al., 2021). The degradation and loss of forests is accelerating in many regions of the world due to global climate change. Assessing this loss in forest cover is relatively straightforward with remote sensing tools, and the consequences on net global carbon balance and other biogeochemical processes can be modeled (e.g., Bondeau et al., 2007). However, estimating the effects on species diversity and interactions remains challenging, especially for the canopy community, which is difficult to access. To address this knowledge gap, reliable data on species occurrence are needed across spatial and temporal scales, as well as across different trophic levels (Seibold et al., 2018 a). However, this is difficult to achieve with classical canopy habitat assessment methods.

Ozanne (2005) reviewed established techniques and methods to sample and assess canopy arthropods. The simplest technique is to access the canopy directly via rope climbing, canopy walkways, or canopy cranes to collect samples. However, these methods require experience in tree climbing or setting up permanent platforms and walkways (Parker et al., 1992). Further commonly applied techniques are based on chemical knockdown using insecticides (Leroy et al., 2022). The chemicals are distributed through fogging (i.e., hot clouds of chemical droplets rising upwards) or mist blowing (i.e., blowing an air current with chemical droplets into the canopy). The stunned, falling insects are caught in collection hoops and can be identified by morphological assessment (Pedley et al., 2016) or bulk sample DNA metabarcoding (Creedy et al., 2019). Another approach is branch bagging and clipping (i.e., covering the branch in a cloth or bag and cutting the branch), which has the advantage of directly correlating species richness or density with plant or leaf biomass (Krehenwinkel et al., unpublished data). Trapping methods with defined entry areas, such as canopy malaise traps (Skvarla et al., 2021) and flight interception traps (Kowalski et al., 2011), have also been applied to record canopy insects, and vertically stratified artificial substrates have been employed to sample and analyze the distribution of arthropods such as deadwood beetles (Seibold et al., 2018 b).

However, none of these canopy invertebrate diversity monitoring methods are fast, taxonomically comprehensive, noninvasive, and simultaneously cost-efficient. Herein, we propose and test a potential complementary canopy invertebrate monitoring technique involving the collection of rainwash environmental DNA (eDNA). Crucially, eDNA can be extracted from environmental samples such as soil, water, or air, without first isolating any target organisms (Taberlet et al., 2012). Today, eDNA metabarcoding is an established method in marine, freshwater, and terrestrial biodiversity research (Deiner et al., 2017). However, the potential of eDNA metabarcoding of rainwater to assess canopy insect diversity has not been explored. Valentin et al. (2021) investigated the effect of rain on the fate of arthropod eDNA and found that rainfall or mist removes most terrestrial eDNA present on vegetation surfaces. Building on this idea, we performed a simple proof-of-principle analysis and hypothesized that (i) using a rainwash sampler, canopy invertebrates can be detected reliably by eDNA metabarcoding of the collected water shortly after a rain event, and that (ii) the taxonomic composition of collected communities will differ between the tree host taxa, with distinct host-specific invertebrates being found below different canopies.

## Methods

### Rain sampler

Four rain samplers were built using 1 m^2^ of 0.5 mm PVC pond liner (Sika, Stuttgart Germany), eight 1 m PVC tubes with a 50 mm diameter, four PVC three-way tube connectors with a 50 mm diameter (HT CONNECT, Pegnitz, Germany), 20 reusable zip ties, and 22 copper eyelet rings (Vastar, Shanghai, China). First, the pond liner was cut to 1 × 1 m, five holes were punched on all four sides, and copper eyelet rings were used to support the holes (supplementary Figure 1). Two additional holes as overflow outlets were implemented and supported with eyelet rings 25 cm from the center of the liner. This allowed for ∼4 L of water to be collected while protecting the liner from tearing due to weight. The liner was sterilized by applying 1% bleach, which was then washed off using 80% ethanol, followed by deionized water. The liner was then sterilized using UV radiation for 30 min, folded with sterile gloves, and placed in a plastic bag. In the field, four of the 1 m PVC tubes were connected to a square using the three-way tube connectors. The liner was then placed in the frame using sterile gloves and fastened using 20 reusable zip ties while leaving enough room for the liner to expand when water is collected. The other four PVC tubes were then inserted into the three-way connector as legs.

### Sampling sites

The sampling sites were located within a >1000 ha forest area in the lower Rhine region of Germany (N 51.707104, E 6.549781). Four rain samplers were set up the evening before a major rain event on June 19th, 2021 (Figure 1). Two were placed under the canopies of beech (*Fagus sylvatica*, site S1) and oak (*Quercus robur*, S4) broadleaf trees, and two were placed beneath the canopies of larch (*Larix* sp., S2) and pine (*Pinus sylvestris*, S3) coniferous trees. Site S1 was an old-growth beech forest, and S2 was a planted larch monoculture (see Supplementary Figure 2). The sites were between 200 m (S3 and S4) and 3 km (S1 and S3) apart (Figure 2).

**Figure 1:**
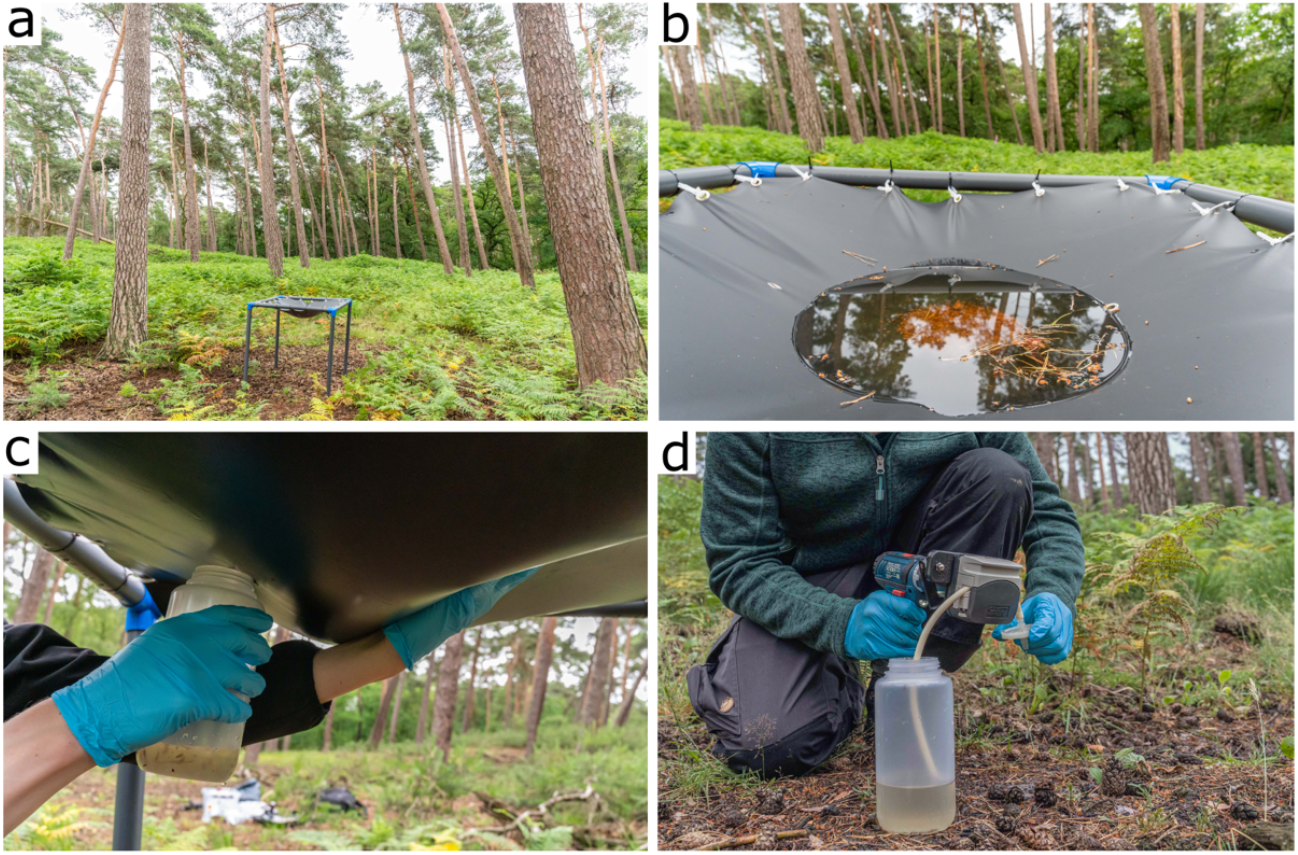
Example of the setup at sampling site S3 in a pine forest (a). Approximately 4 L of rainwater was collected in the liner (b) and was transferred into sterile bottles by pushing the water toward the overflow hole (c). A 2 L volume of rainwater was then filtered using a hand-held peristaltic pump and sterile encapsulated PES filters (d).

**Figure 2:**
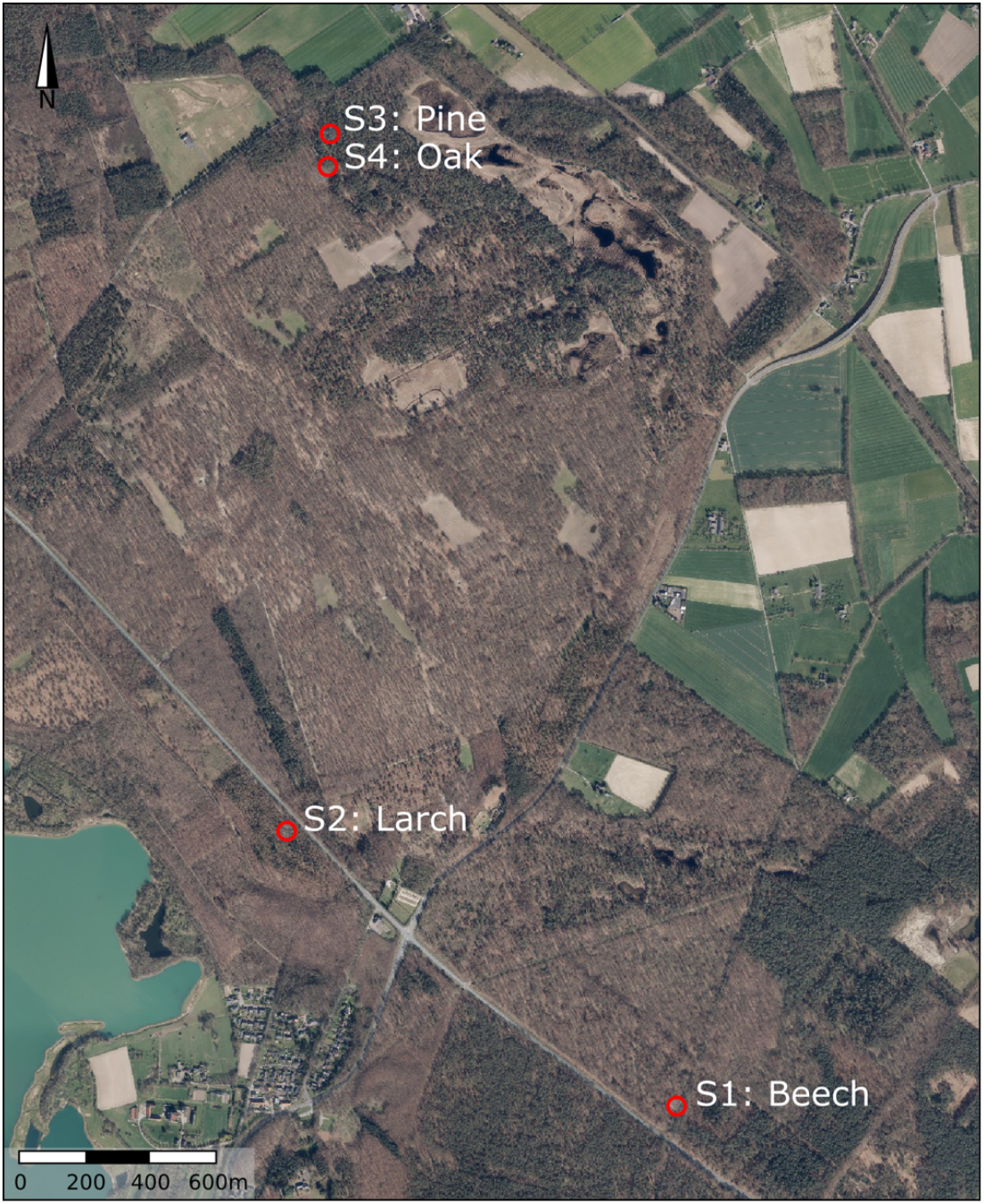
Map of the sampling sites located near Wesel, Germany. Rain samplers were placed below four different tree taxa: beech (*Fagus sylvatica*), larch (*Larix* sp.), pine (*Pinus sylvestris*), and oak (*Quercus robur*). Orthophotos Geobasis NRW, TIM-online 2.

### Verification specimens

To confirm whether the collected eDNA originated from the canopy (washed from leaves and branches) or solely from organisms falling or flying into the rain sampler, all organisms present in the rain sampler were collected. For this, invertebrate specimens were picked using forceps and stored in 80% ethanol. Specimen collection was conducted after eDNA sampling to prevent contamination of the rainwater. All collected specimens were morphologically identified to at least order level using a ZEISS Stemi SV 11 stereo microscope (Oberkochen, Germany).

### eDNA metabarcoding

A detailed description of the eDNA metabarcoding workflow can be found in Supplementary Material 1. Briefly, 2 L of rainwater was collected 45 h after setting up the samplers. The water was immediately filtered on site next to the rain sampler using a Vampire Sampler peristaltic pump (Buerkle, Bad Bellingen, Germany) and collecting eDNA on Whatman Polydisc AS disk filters (PES, 50 mm diameter, 0.45 μm pore size, sterile, Maidstone, UK). Subsequently, DNA was extracted in a sterile lab using an adapted NucleoMag tissue kit (Macherey Nagel, Düren, Germany). A two-step PCR approach was applied to amplify the extracted DNA, with fwhF2 and fwhR2n invertebrate primer pair (Vamos et al., 2017), which is known to reliably amplify DNA from terrestrial insects (Elbrecht et al., 2019). Samples were sequenced on an Illumina HiSeq 2×150 platform at Macrogen (Seoul, Rep. of Korea). Raw reads were received as demultiplexed fastq files. All samples were processed with the APSCALE-GUI pipeline v1.2.0 (Macher et al., unpublished data). Taxonomy was assigned using BOLDigger (Buchner & Leese, 2020). The resulting taxonomy and read table were then converted to a TaXon table and filtered prior to downstream analyses in TaxonTableTools v1.4.1 (TTT, Macher et al., 2021).

### Data analysis

To assess if the rainwash eDNA data differed between the four sites/host tree taxa, we analyzed relative read abundances per invertebrate species across the four host tree taxa. Pragmatically, species with ≥70% relative read abundance to one of the four host tree taxa were classified as ‘beech,’ ‘oak,’ ‘larch,’ ‘pine,’ or ‘undefined.’ The same was done for operational taxonomic units of fungi (OTUs; Ascomycota and Basidiomycota). To assess the host specificity of phytophagous insects, feeding type preference was assigned to species in the eDNA taxa list (‘oak,’ ‘larch,’ ‘pine,’ or ‘broad-leaved trees’), and occurrence patterns on host species were analyzed based on relative read abundances.

To investigate whether the species detected by eDNA metabarcoding were true signals from the canopy above the samplers (i.e., rainwash eDNA) or limited to signals derived from specimens that fell into the rain sampler during sample collection (verification specimens), both species lists were compared.

## Results

### eDNA metabarcoding

Sequencing yielded a total of 30,005,824 raw reads. In total, 10,631,775 quality-filtered reads were clustered into 982 OTUs. While the average number of reads per sample was 664,485 reads (± 197,104), 647 reads (± 315) were assigned to field blanks and negative controls (8 OTUs). After PCR replicate merging and subtraction of the sum of reads per OTU that were present in the field blanks and negative controls, 389 OTUs with similarity ≥85% to reference sequences remained. For downstream analyses, we kept only OTUs of the phyla Arthropoda (48 assigned species), Tardigrada (2), and Nematoda (0). Most species belonged to the orders Lepidoptera (17) and Coleoptera (13), while the remaining 17 orders were represented by fewer than three species each (Supplementary Figure 3).

In total, we detected 21 species under both broadleaf trees (beech and oak) and conifers (larch and pine), while 20 and 9 species were exclusively found under broadleaf trees and conifers, respectively. When using relative read abundances as a proxy, more distinct occurrence patterns were observed; 20 species were detected mostly under oak, while eight species were mainly found under beech (Figure 3). For conifers, nine species were found mainly under larch, and seven species mainly beneath pine. The remaining six species showed no distinct occurrence patterns toward any specific tree taxa. However, four of the taxa were predominantly found under conifer trees.

**Figure 3:**
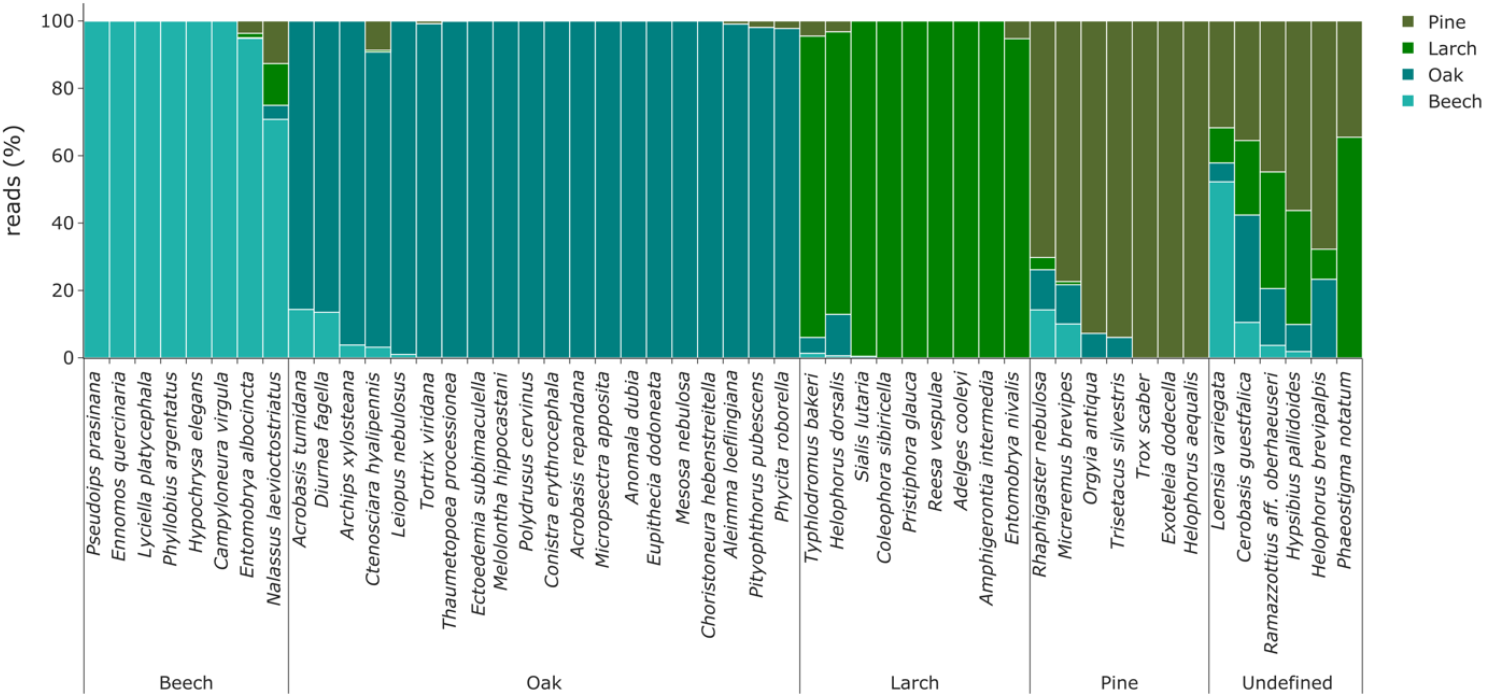
Relative read abundances per site for all detected invertebrate species. Species are grouped by tree taxa according to their relative read abundance (threshold ≥70% relative read abundance).

In total, 25 phytophagous species were detected in the eDNA data. Of these, five species specialized on oak, compared with two on pine and two on larch. A further 16 species are described with linkages to broad-leaved trees, while the habitat of one phytophagous species is not specified. In 21 cases, the linkage was congruent with the species occurrence patterns observed in the eDNA results (Figure 4). For three species, detection was less congruent with their ecology.

**Figure 4:**
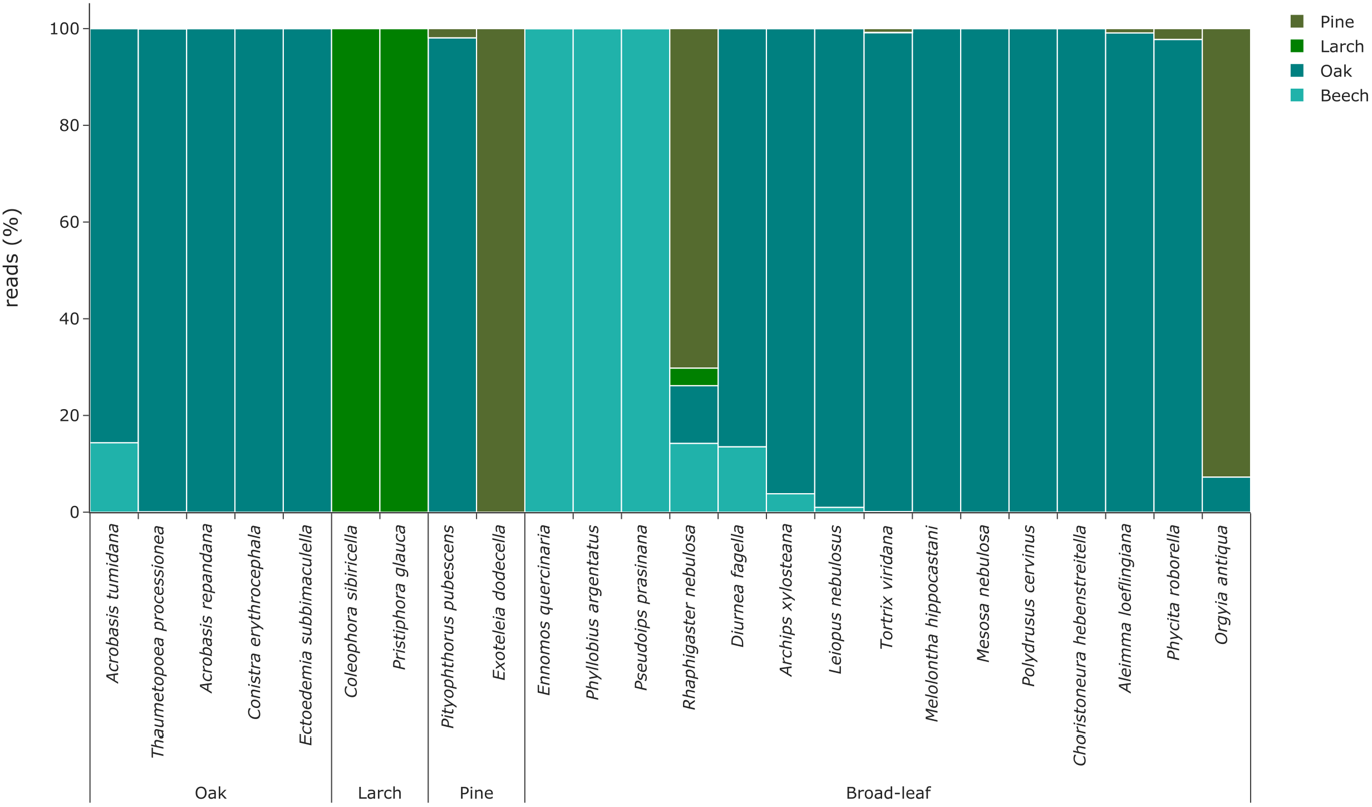
Relative read abundances of phytophagous insects according to the four host tree taxa studied.

### Verification specimens

We sampled 220 invertebrate specimens from the rain samplers after eDNA filtration and identified 18 invertebrate taxa (13 species) based on morphological identification. The specimens belonged to the orders Coleoptera (4 species), Hymenoptera (4), Diptera (1), Hemiptera (1), Isopoda (1), Julida (1), and Lepidoptera (1). When comparing the identified species that were present in the water to the species detected by eDNA analysis, 43 species were detected only by eDNA, seven species were observed with both methods, and six species were only detected through morphological identification (Figure 5).

**Figure 5:**
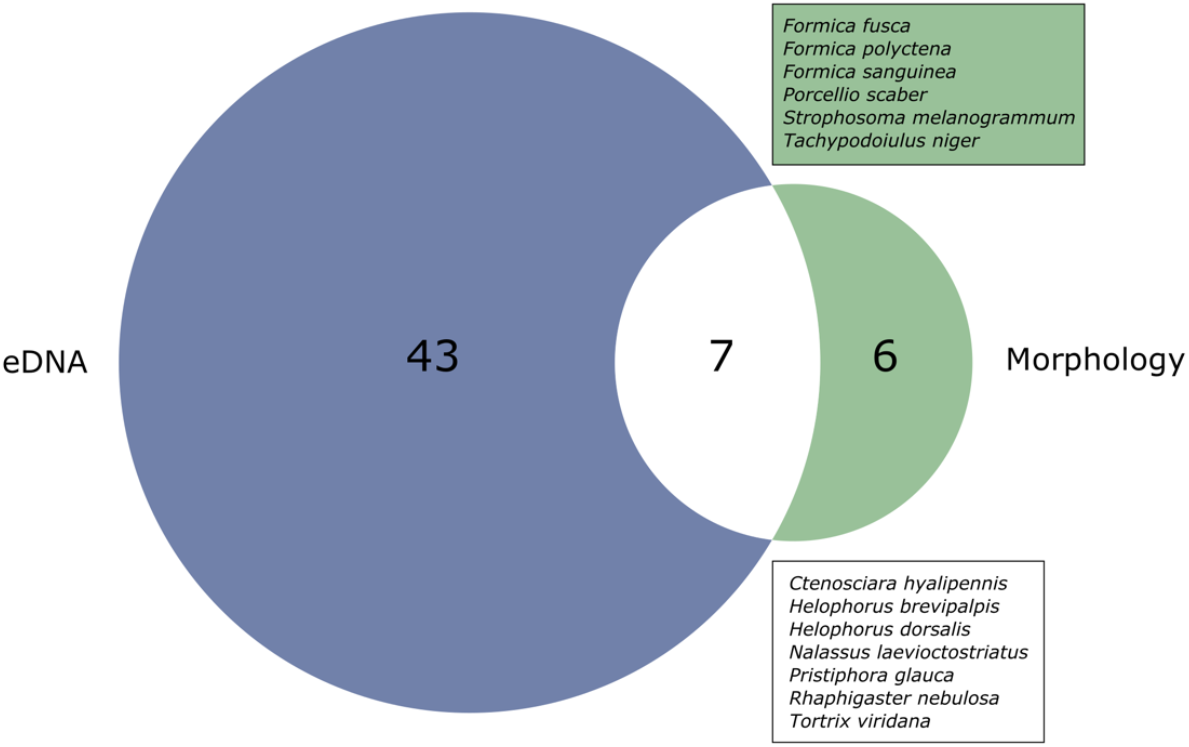
Number of species that were detected by eDNA metabarcoding alone (blue), by both methods (white box), or exclusively by morphological identification of specimens collected from the rain sampler (green box).

## Discussion

Our results support the first hypothesis that eDNA metabarcoding of rainwater collected below the tree canopy can detect many canopy invertebrate species. Despite the small number of samples collected in our pilot study, we detected a remarkable number of invertebrate species from rainwater eDNA. To test whether the eDNA signals were derived from eDNA sources in the canopy, we also collected and identified all specimens that fell into the rain sampler to verify the eDNA signals. We found significantly more species through eDNA metabarcoding than through morphological identification. Additionally, all species exclusively identified through morphological identification were present in low abundances. Three of the six species were small ants (Formicidae), which most likely do not shed large amounts of eDNA into the water. The detected aquatic beetles of the genus *Helophorus* likely colonized the small water bodies created in the rain samplers as they usually inhabit puddles. Overall, these results prove that most species detected within the rainwash water were eDNA signals washed into the sampler from the tree canopy. This aligns with results reported by Valentin et al. (2021) showing that invertebrate eDNA is washed off vegetation surfaces and can be detected. However, it is highly likely that only a very small proportion of the eDNA released by the invertebrate community in the canopy was detected in rainwash eDNA, since the samplers only covered an area of 1 m^2^. Additionally, the observed species richness will most likely vary considerably with different rain intensities. The minimal amount of precipitation needed to recover most of the canopy community via rainwater metabarcoding needs to be explored in future studies. Nevertheless, our results revealed an impressively large number of canopy invertebrate species in the rainwater. By comparison, Leroy et al. (2022) found 757 putative species by sampling 40 trees in oak-dominated stands, using canopy fogging and DNA metabarcoding. This proves that even a small subsample of the whole rainwater that passes through the canopy can offer substantial and novel insights into the invertebrate community. To further maximize detection, more samplers covering a greater area would be needed.

Our results also support our second hypothesis that eDNA can reveal differences in taxonomic composition between different tree species. Despite the limited number of rain samplers used in this study, and hence the limited tree canopy coverage, the rainwash eDNA results revealed distinct species occurrence patterns under the four different trees. In total, 88% of detected species were assigned to a certain tree taxon based on their relative read abundances. In fact, many of those species have a known host specificity toward specific tree taxa, such as *Acrobasis repandana* to oak, *Exoteleia dodecella* to pine, and *Pristiphora glauca* to larch (Supplementary Material 1). Other species, such as *Trox scaber*, occupy bird nests and have only a secondary ecological linkage to trees (Supplementary Material 1). Additionally, our unfiltered data included a remarkable number of fungal OTUs displaying tree-specific occurrence patterns (Supplementary Figure 4). This provides evidence that fungal eDNA was washed from the canopy alongside invertebrate eDNA, and that both can be extracted from rainwater, which opens the possibility for multimarker analyses with additional fungi-specific DNA metabarcoding markers. This facilitates the analysis of even more complex multitrophic ecological interactions, such as invertebrate and fungi co-occurrence patterns.

Our results demonstrate the potential of rainwash eDNA metabarcoding as a rapid and minimally invasive method for measuring canopy invertebrate diversity. While our occurrence data have limited statistical power, due to the small sample size, they suggest that local canopy communities can be distinguished using rainwash eDNA. In particular, phytophagous insects that specialize on single host tree species were detected locally in our results, which demonstrates the potential for applications in forestry or forest sciences.

To generate statistically robust results, future studies with more comprehensive designs are required. For example, several rain samplers should be set up per forest type, and several forests of the same type could be investigated. For this, stationary and passive hard-shell rain samplers could be implemented in forest survey areas, with regular emptying. To record forest-specific communities or target communities of specific shrub or tree species, rain samplers could be installed at different heights in the canopy. In urban setups, rain samplers could potentially be installed in water catchment trays that are often used to enhance the supply of water to urban trees. Specific collection of metadata (e.g., tree height, diameter at breast height, crown density) could also generate multivariate data in addition to the samples and could be used for a more standardized setup. Since our rainwash eDNA metabarcoding approach relies on natural rain events, its most promising field of application lies in the canopy biomonitoring of rainforests or areas with regular precipitation. However, in drier regions, actively rinsing eDNA off bushes or tree canopies with a water hose could be an alternative approach, as already conducted by Valentin et al. (2020) for species-specific assessments.

In conclusion, rainwash eDNA metabarcoding has the potential to substantially advance forest canopy biodiversity monitoring. Our results highlight the possibility of a minimally invasive, cheap, and comprehensive approach, which could even be expanded to complex multitrophic analyses. With further improvements, our method could significantly contribute to closing the gaps in our knowledge of biodiversity and ecological interactions of canopy communities.

## Acknowledgments

We would like to thank the Lower Landscape Authority of the District of Wesel and the Landesbetrieb Wald und Holz NRW for enabling our investigations. We thank members of the Leese lab for comments and feedback.

## Author contributions

RS, TM, TH, AB, and FL conceived the study design. RS and TM conducted eDNA sampling. TH collected and identified verification specimens and provided expertise on traits. TM conducted the laboratory workflow and bioinformatic analyses. TM and RS drafted the first version of the manuscript. All authors contributed to the article and approved the version to be published.

## Conflict of interest

The authors have no conflicts of interest to declare.

## Supplementary figures

**Supplementary Figure 1:**
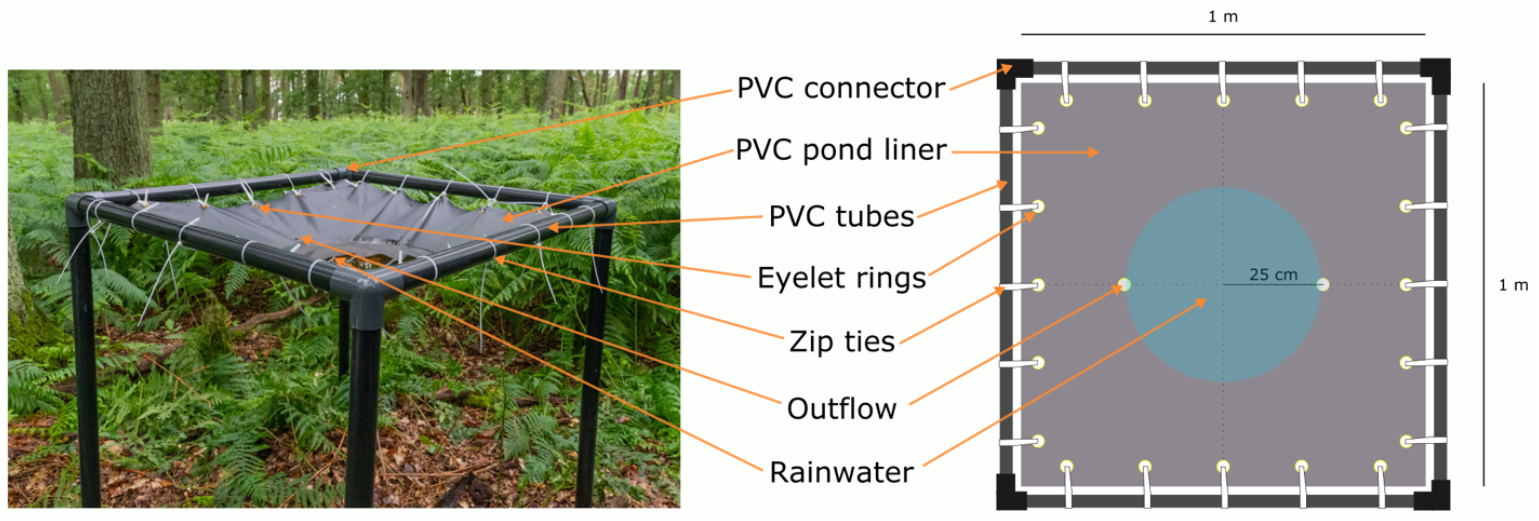
Schematic overview of the rain sampler prototype.

**Supplementary Figure 2:**
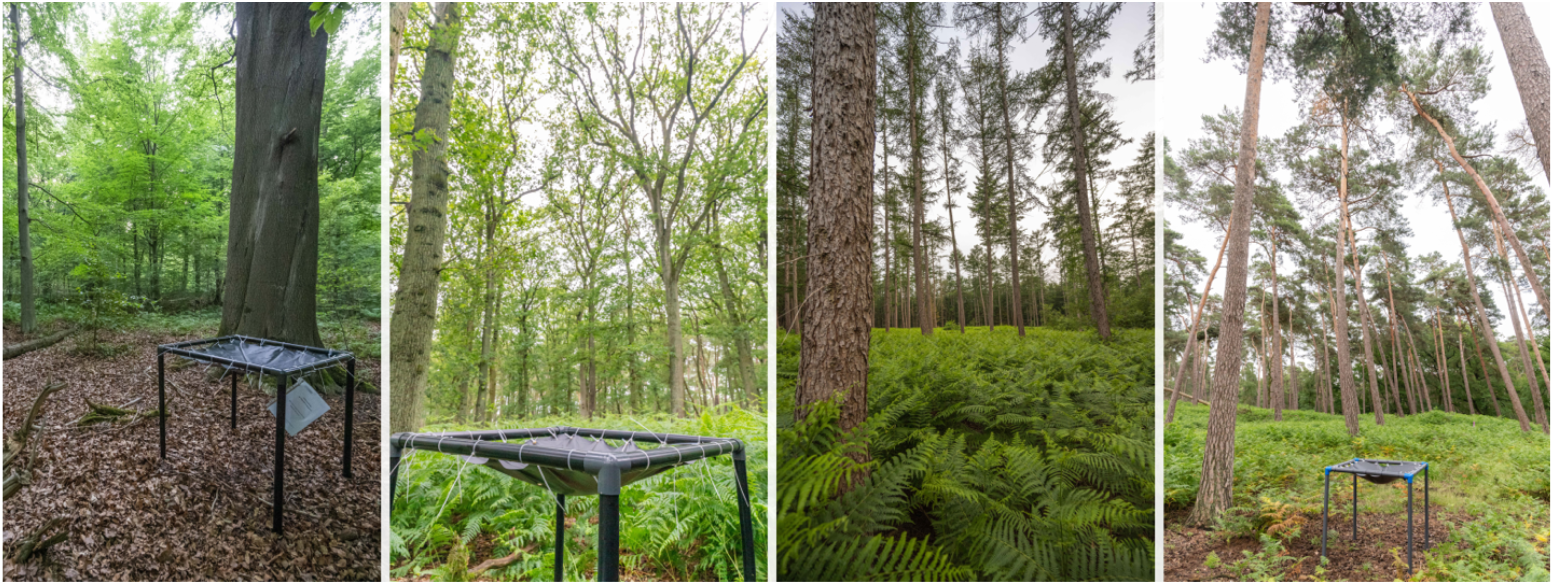
Sampling sites under the four different tree taxa (from left to right: S1 beech, S4 oak, S2 larch, S3 pine).

**Supplementary Figure 3:**
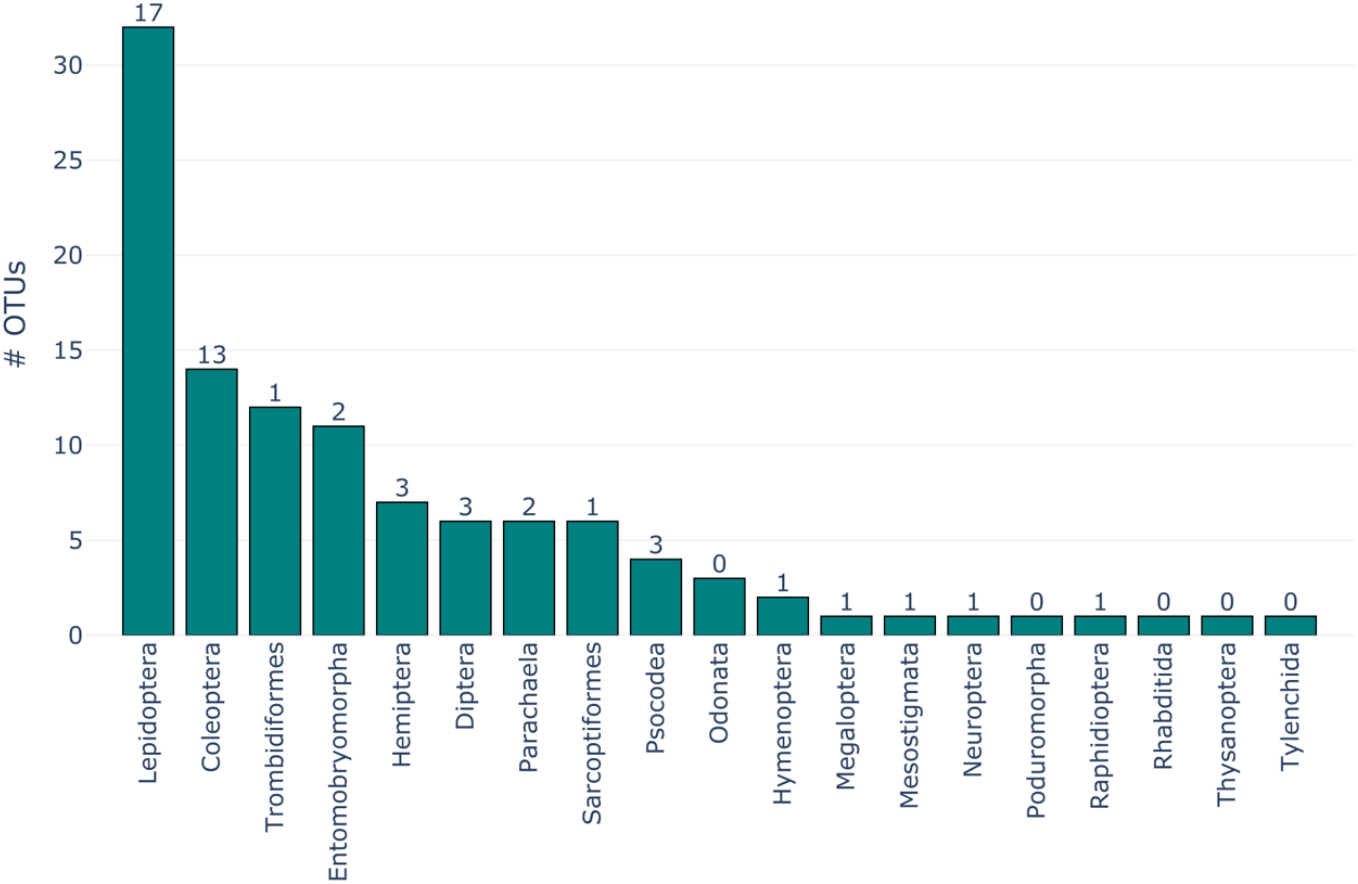
Number of invertebrate OTUs per order of the rainwash eDNA dataset. The number of detected species is shown above the respective bar.

**Supplementary Figure 4:**
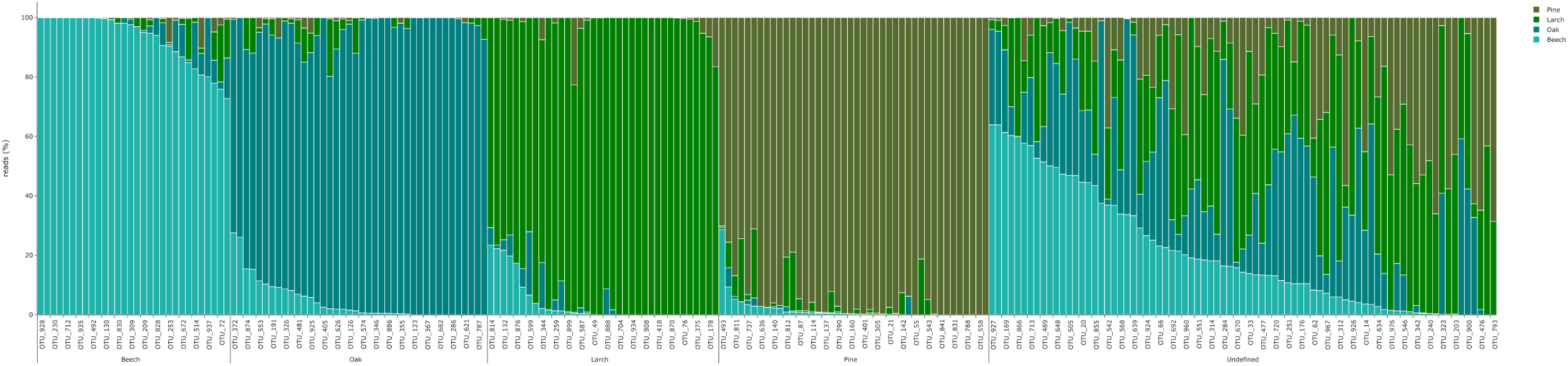
Relative read abundances of fungi OTUs (Ascomycota and Basidiomycota) per site (Beech, Oak, Pine, Larch). OTUs are grouped by tree taxa according to their relative read abundance (threshold ≥70% relative read abundance).

## Notes

### Competing Interest Statement

The authors have declared no competing interest.

