## Supplementary Material 1 for "It’s raining species: Rainwash eDNA metabarcoding as a minimally invasive method to assess tree canopy invertebrate diversity"

### Material and methods

#### Sampling sites

The sampling sites were located within a >1000 ha forest area in the lower Rhine region of Germany (N 51.707104, E 6.549781), which includes the ‘Diersfordter Wald’ nature protection area and the ‘Großes Veen’ EU Special Area of Conservation (92/43/EEC). Multiple different forest types with different tree species occur in this area. Four rain samplers were set up the evening before a major rain event on June 19$^{th}$, 2021. Two were placed under the canopies of beech (*Fagus sylvatica*, site S1) and oak (*Quercus robur*, S4) broadleaf trees, and two were placed beneath the canopies of larch (*Larix* sp., S2) and pine (*Pinus sylvestris*, S3) coniferous trees. Site S1 was an old-growth beech forest, and S2 was a planted larch monoculture (see Supplementary Figure 3). The sites were between 200 m (S3 and S4) and 3 km (S1 and S3) apart (Figure 1).

#### eDNA sampling

A 2 L volume of water was collected from each rain sampler in a sterile 1 L bottle (Nalgene) 45 h after setting up the samplers. Within this timeframe, the closest weather station (20 km distance) reported a combined precipitation of 35 mm, while the temperatures were between 16°C at night and 26°C during the day (data from wetteronline.de for Duisburg-Baerl, June 2021). Two individual bottles per site were filled with water that was discharged through the overflow outlet in the pond liner. The water was immediately filtered on site next to the rain sampler by pumping the water from the bottle using a Vampire Sampler peristaltic pump (Buerkle, Bad Bellingen, Germany) and collecting eDNA using Whatman Polydisc AS disk filters (PES, 50 mm diameter, 0.45 µm pore size, sterile, Maidstone, UK). After filtration, TNES buffer (50 mM Tris, 0.4 M NaCl, 100 mM EDTA, 0.5% SDS) was added to the disk filters using 1 mL syringes (Braun, Melsungen, Germany), and filters were closed using sterile rubber plugs. The filters were stored on ice in the field and placed at -20°C in a freezer until extraction.

**eDNA extraction**

All wet lab steps were conducted under sterile conditions in a dedicated sterile laboratory (UV lights, sterile benches, overalls, gloves, and face masks). First, 100 µL TNES buffer and 10 µL Proteinase K (7BioScience, Neuenburg am Rhein, Germany) were added to the inlet of each filter, and all filters were incubated at 55°C with shaking at 1000 rpm for 3 h on an Eppendorf ThermoMixer C instrument (Eppendorf AG, Hamburg, Germany). After lysis, liquid was extracted from the filter using a sterile 5 mL syringe (Braun, Melsungen, Germany) and transferred to a new 2 mL Eppendorf tube. Subsequently, DNA was extracted using an adapted NucleoMag tissue kit (Macherey Nagel, Düren, Germany; Supplementary Material 1). In total, a volume of 400 µL per sample was extracted.

**eDNA amplification and sequencing**

A two-step PCR approach was applied for amplifying the extracted DNA. In the first PCR step, tagged versions of primers fwhF2 and fwhR2n (Vamos et al. 2017) were used, which are optimized for invertebrates, target a 205 bp cytochrome c oxidase subunit 1 (COI) gene fragment, and are known to reliably amplify DNA from terrestrial insects (Elbrecht et al., 2018). In total, 19 first-step PCR amplifications were conducted, including two replicates per sample, two negative PCR controls, and one field blank. The reaction volume was 25 µL, consisting of 7 µL PCR-grade water, 12.5 µL Multiplex Mastermix (Qiagen Multiplex PCR Plus Kit, Qiagen, Hilden, Germany), 0.5 µL fwhF2 forward primer, 0.5 µL fwhR2n reverse primer, and 5 µL of DNA template. The first

PCR step was carried out at 95°C for 5 min, followed by ten touchdown cycles at 94°C for 30 s, 68–59°C for 90 s, and 72°C for 30 s, followed by 25 cycles at 94°C for 30 s, 58°C for 90 s, and 72°C for 30 s. The final elongation was carried out at 68°C for 10 min. In the second PCR step, Illumina sequence adapters with a combinatorial indexing system were added (Buchner et al., 2021). For each sample, the second-step PCR mix contained 7 µL PCR-grade water, 12.5 µL Multiplex Mix, 1 µL combined primer (10 µM), and 5 µL first-step product. PCR conditions were 95°C for 5 min, followed by 15 cycles at 94°C for 30 s and 72°C for 120 s. The final elongation was carried out at 68°C for 10 min. Following second-step PCR, products were visualized on a 1% agarose gel to evaluate amplification success. Samples were subsequently normalized to 25 ng per sample using a SequalPrep Normalization Plate (Applied Biosystems, Foster City, CA, USA) following the manufacturer's protocol. Subsequently, normalized samples were pooled into one library with samples of a different project. The pooled library was concentrated using a NucleoSpin Gel and PCR Clean-up kit (Macherey Nagel) following the manufacturer's protocol. The final elution volume of the library was 40 µL. The library was then analyzed using a Fragment Analyzer (High Sensitivity NGS Fragment Analysis Kit; Advanced Analytical, Ankeny, USA) to check for potential primer dimers and coamplification, and to quantify the DNA concentration of the library. Primer dimers were removed by cutting the library from a 2% agarose gel and extracting the DNA from the gel using a NucleoSpin Gel and PCR Clean-up kit (Macherey Nagel). Subsequently, samples were normalized and pooled, and the final library was concentrated. The resulting library (together with samples from a different project) was sequenced on a HiSeq 150 bp PE Illumina platform at Macrogen (Seoul, Rep. of Korea).

**Bioinformatics**

Raw reads were received as demultiplexed fastq files. All samples were processed with the APSCALE-GUI pipeline v1.2.0 (Macher et al., 2022 under review), which is based on VSEARCH (Rognes et al., 2016) and cutadapt (Martin, 2011). All settings were kept as default, and OTUs were clustered with a 97% percentage similarity threshold. Subsequently, taxonomy was assigned using BOLDigger (Buchner & Leese, 2020), which automatically performs an identification search against the BOLDsystems COI database (www.boldsystems.org). The resulting taxonomy table was filtered using the 'JAMP filtering' option. Both the taxonomy and read table were then converted to a TaXon table (Supplementary Material 2) for downstream analyses in TaxonTableTools v1.4.1 (TTT, Macher et al., 2021).

Initially, PCR replicates were merged, and only OTUs that were present in both PCR replicates were kept, and the sum of reads of all OTUs that were present in negative controls were subtracted from the samples. The dataset was then filtered by taxonomic groups, and only OTUs with similarity >85% to the nearest reference sequence and that were assigned to the phyla Arthropoda or Tardigrada were kept. This TaXon table was used for all downstream analyses (Supplementary Material 3). Additionally, a taxon list was created in which OTUs with the same taxonomy were merged to a single entry (Supplementary Material 4).

Despite read abundances being distorted by factors such as differences in binding efficiencies or DNA shedding rates, they hold some quantitative information (Krehenwinkel et al., 2017). Thus, relative read abundances per invertebrate species were analyzed for distribution patterns across the four tree taxa. Therefore, species with ≥70% relative read abundance to one of the four tree taxa were categorized as 'beech,' 'oak,' 'larch,' 'pine,' and 'others.' Occurrence patterns were plotted as a bar chart. The distributions of fungi OTUs (Ascomycota and Basidiomycota) were investigated accordingly.

Furthermore, ecological traits were added to the taxon list (Supplementary Material 4). Therefore, species were categorized according to their phytophagous larvae ecology (oak, larch, pine, or broadleaved trees). To assess the larva ecology linkage based on the eDNA metabarcoding results, the detected species were categorized by their respective traits and occurrence patterns (relative read abundance) and plotted as a bar chart.

To investigate whether the species detected by eDNA metabarcoding were true signals from the canopy above the samplers (i.e., rainwash eDNA) or limited to signals derived from specimens that fell into the rain sampler during sample collection (verification specimens), both species lists were compared and visualized in a Venn diagram.

**Supplementary figures**

Supplementary Figure 1: Schematic overview of the rain sampler prototype.

Supplementary Figure 2: Sampling sites under the four different tree taxa (from left to right: S1 beech, S4 oak, S2 larch, S3 pine).

Supplementary Figure 3: Number of invertebrate OTUs per order of the rainwash eDNA dataset. The number of detected species is shown above the respective bar.

Supplementary Figure 4: Relative read abundances of fungi OTUs (Ascomycota and Basidiomycota) per site (Beech, Oak, Pine, Larch). OTUs are grouped by tree taxa according to their relative read abundance (threshold ≥70% relative read abundance).
